# PHYTOCHROME INTERACTING FACTORS are required to coordinate microtubule dynamics and differential cell growth during Arabidopsis apical hook opening

**DOI:** 10.1101/2025.02.18.638861

**Authors:** Natalia Burachik, Paula Vacs, Santin Franco, María Agustina Mazzella, Nahuel González-Schain

**Affiliations:** Instituto de Investigaciones en Ingeniería y Biología Molecular “Dr. Hector N. Torres” (INGEBI); CONICET. Buenos Aires, Argentina; Instituto de Biología Molecular y Celular de Rosario (IBR); CONICET-UNR. Rosario, Argentina

## Abstract

**Background:** Differential cell growth is a key strategy that enables plants to adjust their morphology in response to internal and external cues. Apical hook opening in dark-grown Arabidopsis seedlings serves as an excellent model to study how coordinated cell expansion generates organ bending. Several works highlight the influence of hormone signaling, cell wall mechanics, and environmental cues on cortical microtubule (cMT) arrangement in bending tissues. However, the precise genetic hierarchy governing these processes remains largely unknown.

**Results:** Here, we reveal a two-step growth rate during apical hook opening, in which an initial phase of differential cell expansion is followed by a phase of more balanced growth. We show that cMT re-arrangement precedes and predicts cell expansion patterns, suggesting a central role in coordinating differential growth. Our findings indicate that PIF transcription factors regulate this process, as *pifq* mutants fail to align cMT reorganization with growth dynamics, resulting in premature hook opening. RNA-seq analysis further supports a role for PIFs in coordinating cell wall remodeling and microtubule-associated gene expression, including kinesins and cell wall-modifying enzymes.

**Conclusions:** Our study uncovers a PIF-dependent regulatory mechanism that orchestrates cytoskeletal dynamics and cell expansion to control apical hook opening.

## INTRODUCTION

Plants exhibit a remarkable plasticity to adjust their growth in response to environmental cues. Differential cell growth, an evolutionary key strategy, involves unequal cell expansion to bend organs like stems or roots, enabling directional growth. This targeted response helps the plant to adjust growth direction to optimize access to resources and avoid harmful conditions. Apical hook development in dark-germinated seeds is an excellent model to study differential cell growth and organ bending. After germination seedlings rapidly elongate their hypocotyls to ensure they reach sunlight before exhausting embryo reserves, while the development of an apical hook helps plants navigate through the soil by protecting delicate shoot meristem. In this process, cells on the inner (concave) side of the hook expand less than those on the outer (convex) side, resulting in hypocotyl bending. Apical hook development shows three distinctive phases: the formation phase (within the first day after germination), the maintenance phase, for approximately another day, and the opening phase (Wang *et al*., 2023). Both hook formation and opening imply differential cell growth although regulatory mechanisms involved in this process are not fully shared (Mazzella *et al*., 2014; Wang *et al*., 2023).

Cortical microtubules (cMTs) are pivotal for orchestrating cell growth anisotropy, acting as key determinants of cell shape. By guiding the deposition of cellulose microfibrils, transversely oriented cMTs facilitate cell elongation. Conversely, longitudinally aligned cMTs restrict growth, ensuring controlled expansion. While cMTs alignment has been extensively studied in hypocotyl cells (Ehrhardt and Shaw, 2006), our understanding of how coordinated cMT dynamics between neighboring cells contribute to tissue bending remains incomplete. Apical hook formation provides a compelling example of cMTs’ role in tissue curvature (Baral *et al*., 2021; Peng *et al*., 2022). Reorganization of cMTs is crucial for this process, with the microtubule-associated protein WDL4 playing a critical role in promoting hook opening (Deng *et al*., 2021). During light-induced hook opening, cMTs reorient from a transverse to a longitudinal arrangement, with a faster reorientation rate observed on the convex side (Walia *et al*., 2024). While these works highlight the influence of hormone signaling, cell wall mechanics, and environmental cues on cMT arrangement in bending tissues, the precise genetic hierarchy governing these processes remains largely unknown.

Here we used Arabidopsis apical hook opening to study cMT dynamics. Our findings reveal a two-step cMT reorganization process closely linked to differential growth patterns in convex and concave hook sides: initially a combination of moderate growth in the concave side and longitudinal cMT alignment in convex cells restricts growth, followed by transverse reorientation as growth is balanced between both sides. Furthermore, we demonstrated that the dynamics of cMT rearrangement precede cell growth behavior, a process dependent on the master skotomorphogenic regulators PHYTOCHROME INTERACTING FACTORS (PIFs). In a quadruple PIF mutant *pifq* (*pif1 pif3 pif4 pif5*) this coordination is lost-cells elongate despite maintaining longitudinal cMT arrays-. RNA-seq reanalysis revealed reduced cell wall gene expression and upregulated cMT remodeling genes. Our results suggest that PIFs orchestrate cMT organization and cell elongation, ensuring controlled differential growth during hook opening.

## RESULTS AND DISCUSSION

### Microtubule arrangement patterns predict cell expansion in apical hook opening

While previous studies have focused on static cMT arrangements, exploring their dynamics would shed light in understanding how cMTs actively respond to and integrate mechanical and hormonal signals over time to support differential growth. This dynamic perspective is crucial to uncover the temporal regulation and feedback mechanisms that drive hook opening in darkness, providing deeper insights than static snapshots of cMT organization. To examine the interplay between cMT dynamics and apical hook opening during skotomorphogenesis, we analyzed apical hook angle, cell elongation, and cMT organization in dark-grown seedlings over a two- to four-day period (dD, days in darkness). We used a transgenic line expressing the p35S::GFP-MBD fusion (Marc *et al*., 1998) to visualize cMT arrangement in epidermal cells on confocal images (Fig. 1a).

**Figure 1.**
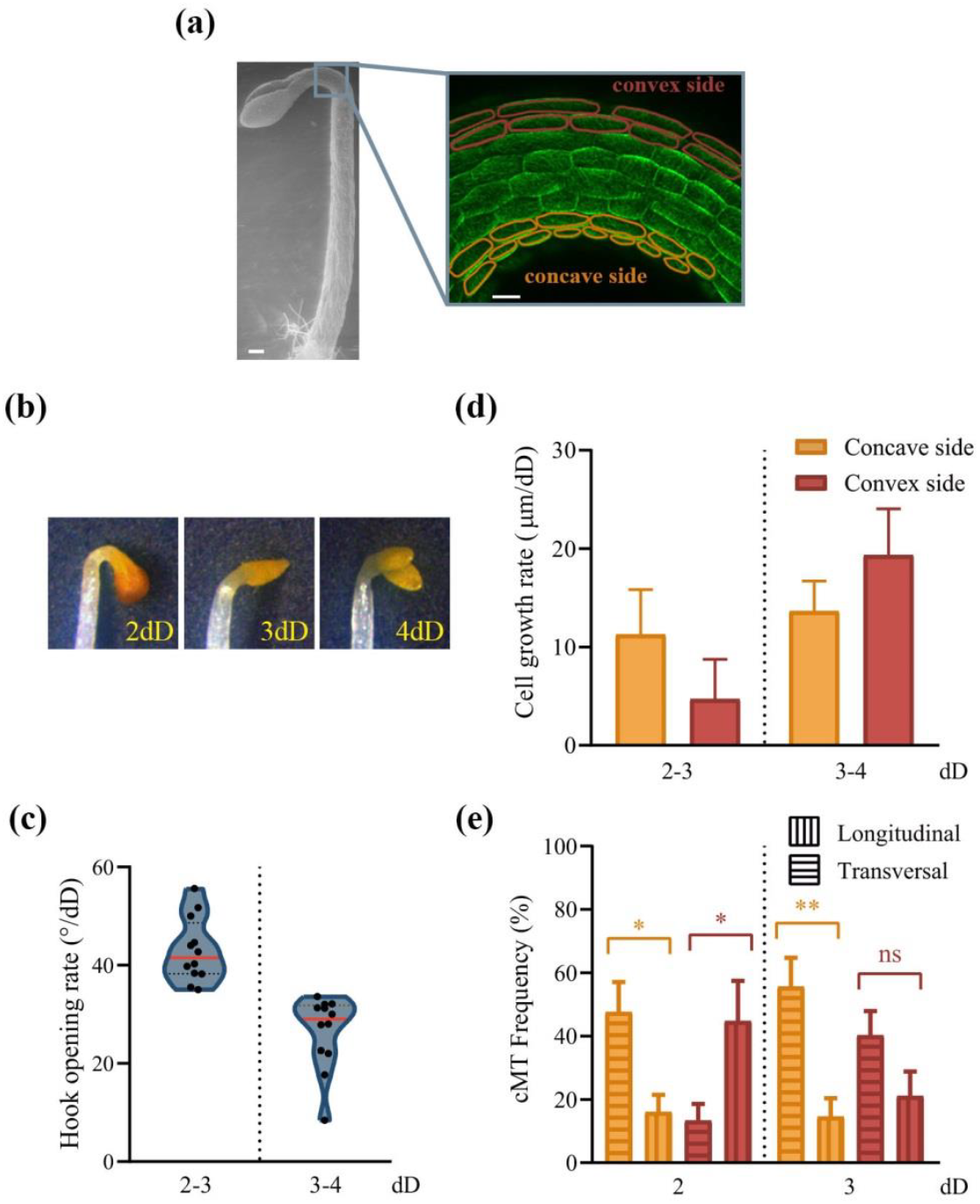
Cortical microtubule re-arrangement during apical hook opening. (a) Scanning electron microscopy image of two-day dark-grown p35S::GFP-MBD cMT marker line (left, bar=100 μm) and confocal fluorescence microscopy image of zoomed hook cells (right, bar=20 μm). (b) Representative p35S::GFP-MBD hooks of seedlings grown in darkness. dD indicates days in the dark. (c) Hook opening rate of p35S::GFP-MBD seedlings between indicated time points. Data are reported as violin plots with median in red, and points represent biological replicates of 30-52 seedlings each. Associated data with hook angle measurements for each time point is presented in Fig. S1a. (d) Cell growth rate of concave and convex side of p35S::GFP- MBD hooks between indicated time points. Data are reported as means ± SEM of at least five biological replicates (total n° of cells=106-189). Associated data with cell length measurements for each time point is presented in Fig. S1b. (e) Frequency of longitudinal and transversal cMTs in concave and convex side of p35S::GFP-MBD hooks at indicated time points. Data are reported as means ± SEM of at least nine biological replicates (total n° of cells=124-143). Asterisks indicate statistically different mean values (t-test, *, P < 0.05; **, P< 0.01; ns, not significant).

In p35S::GFP-MBD dark-grown seedlings, apical hook opening exhibited two distinct phases: a rapid opening phase between days two and three, followed by a slower phase from day three to four (Figs 1b,c, S1a). This observation aligns with a previous report describing a fast and slow opening stage (Smet *et al*., 2014). Rapid hook opening between days two and three correlated with increased cell elongation on the concave side, while convex side cells exhibited minimal expansion (Figs 1d, S1b). Growth pattern of 2-3dD aligns with the cMT arrangement on day two, where prevalent transverse cMTs on the concave side promote growth, whereas longitudinal cMTs on the convex side restrict growth (Fig. 1e). From day three to four, cell growth accelerated on both sides (Figs 1d, S1b), with hook opening slowing (Fig. 1c). This aligns with the cMTs arrangements on day three, exhibiting a more transverse orientation on both sides (Fig. 1e). These findings indicate that hook opening proceeds through two distinct phases, with cMT orientation preceding cell expansion. During the rapid hook opening phase, cMT alignment supports differential cell growth. Subsequently, an increase in transverse cMTs on the convex side, coinciding with higher cell growth rates, suggests a coordinated shift in cMT dynamics. This shift may contribute to more balanced, slower growth, potentially stabilizing the hook opening process.

The fast hook opening phase observed in this study, occurring between 48 and 72 hours after germination induction, aligns temporally with the period of accelerated growth reported in etiolated hypocotyls in previous research (48 to 60 hours after germination induction) (Refregier *et al*., 2004; Pelletier *et al*., 2010). Under our conditions, hypocotyl growth accelerated between days 2 and 3 and decelerated between days three and four (Fig. S2a,b), mirroring hook opening dynamics (Figs. 1b,c, S1a). Concomitantly, we observed that apical hypocotyl cells also exhibited higher growth rates between days two and three compared to days three and four (Fig. S2c,d). On day two, most cMTs are oriented transversely, a configuration that favors cell expansion and aligns with the increased growth rate during 2-3dD (Fig. S2e). By day three, a larger proportion of cMTs adopt a longitudinal orientation, which limits cell expansion and aligns with the reduced growth rate observed between days three and four (Fig. S2e). Our results suggest a coordinated interplay between cMT organization and cell expansion in apical hypocotyl and hook cells. Despite apical hypocotyl and hook cells belonging to different anatomical structures, our findings raise the intriguing possibility that the observed growth acceleration in apical hypocotyl cells might generate mechanical forces that, together with the coordinated arrangement of cMTs and auxin dynamics (Du *et al*., 2022), contribute to the process of differential cell growth in hook cells leading to hook opening.

### *cMTs arrangement pattern is uncoupled from cell expansion in apical hook opening in* pifq *mutants*

The quadruple mutant *pifq* displays a short hypocotyl and a hook with a slight curvature at 2dD (Leivar *et al*., 2008; Zhang *et al*., 2018; Griffiths *et al*., 2024). Under our conditions, two-day-old dark-grown *pifq* mutants exhibited a nearly fully open apical hook, by 3dD only a microscopic distinction between concave and convex sides was apparent (Fig. 2a), and by 4dD the hook was undetectable. Interestingly, on day two of growth in darkness, cMTs orientation (Fig. 2c) did not correlate with the cell growth rate observed between days two and three (Figs 2b, S3). For instance, despite an ~80 μm increase in cell length on both sides in the *pifq* mutants during this period, there was no predominant transverse cMTs orientation. Moreover, most cMTs on the convex side remained predominantly longitudinally aligned between days two and three (Fig. 2c). Cell growth on the concave side between days three and four was high (Figs 2b, S3), despite cMTs being predominantly arranged longitudinally (Fig. 2c). These results suggest that PIF transcription factors may play a crucial role in coordinating cMTs dynamics and cell elongation in specific cells to facilitate proper hook opening in darkness. Hypocotyls displayed a growth acceleration between two and three days in *pifq* dark-grown seedlings and then completely ceased growth (Fig. S4a,b). Once again, cMT arrangements do not align with cell growth in the apical cells of the hypocotyl in *pifq* mutants (Fig. S4c,e). This misalignment may be linked to the role of PIFs in regulating cMT-related proteins, such as SPIRAL1 (Zhou *et al*., 2023) or the Microtubule-Destabilizing Protein MDP60 (Ma *et al*., 2018).

**Figure 2.**
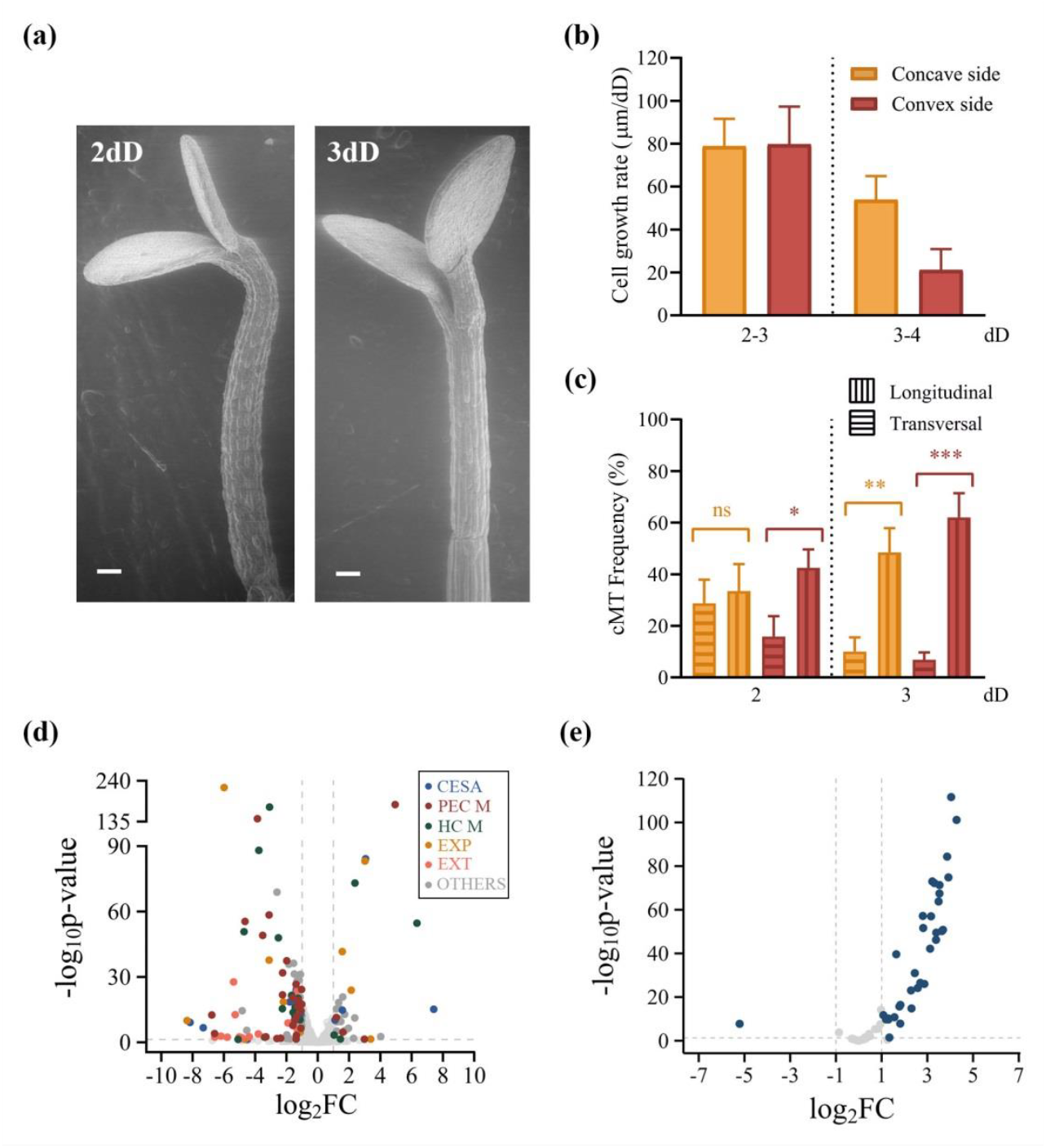
PIF transcription factors coordinate cell wall and cMT homeostasis in apical hooks. (a) Scanning electron microscopy images of two- and three-day dark-grown p35S::GFP-MBD in *pifq* background (bars=100 μm). (b) Cell growth rate of concave and convex side of p35S::GFP-MBD *pifq* hooks between indicated time points. Data are reported as means ± SEM of at least five biological replicates (total n° of cells=60-86). Associated data with cell length measurements for each time point is presented in Fig. S3. (c) Frequency of longitudinal and transversal cMTs in concave and convex side of p35S::GFP-MBD *pifq* hooks at indicated time points. Data are reported as means ± SEM of at least nine biological replicates (total n° of cells=113-121). Asterisks indicate statistically different mean values (t-test, *, P < 0.05; **, P< 0.01; ***, P<0.001; ns, not significant). (d,e) Volcano plots showing log2FC (*pifq*/WT) vs – log10p-value for genes belonging to cell wall organization or biogenesis (GO:0071554) (d), or microtubule motor activity (GO:0003777) (e) from four-day apical hook dark-grown *pifq* and WT RNA-seq re-analysis (Zhang *et al*., 2021). Associated values are presented in Table S1. Horizontal dashed lines represent the p-value threshold, while vertical dashed lines represent cut-off expression values. CESA: cellulose synthase genes; PEC M: pectin metabolism associated genes; HC M: Hemicellulose metabolism associated genes; EXP: expansins; EXT: extensins.

Intriguingly, by day three, cells on both sides of the apical hook in the *pifq* mutant (concave mean ± SEM: 121.5±6.2 μm; convex: 151.6±7.6 μm, Fig. S3) were significantly larger than those in the WT (concave mean ± SEM: 41.5±1.5 μm; convex: 62.7±1.8 μm, Fig S1b), with these nearly threefold size differences persisting at 4dD (Figs S1b, S3). These differences can also be seen in SEM images (Fig. S5). The misregulation of cell expansion, together with the unexpected dynamics of cMTs arrangement observed in *pifq* hooks, prompted us to investigate the underlying factors coordinating these processes. To this end, we re-analyzed an organ-specific RNA-seq data from the apical hook and hypocotyl of wild-type (WT) and *pifq* mutant seedlings at 4dD, using previously reported data (Zhang *et al*. (2021). In the hooks, we observed an enrichment of the Differential expressed genes (DEGs) involved in cell wall organization or biogenesis (Gene Ontology term GO:0071554) (Table S1), consistent with previous report (Zhang *et al*., 2021). These genes encode proteins responsible for the synthesis or modification of cellulose, hemicellulose and pectins (Qiu *et al*., 2021), as well as cell wall modifying proteins like expansins and extensins (Moussu and Ingram, 2023). Notably, approximately one-sixth of the 590 genes in this category were downregulated in *pifq* hooks (Table S1, Fig. 1d), including several cellulose synthase (CESA), xyloglucan endotransglucosylase/hydrolase (XTHs) or xyloglucan xylosyltransferases (XXTs) (grouped within the hemicellulose metabolism genes, HC M, in Fig. 2d), or pectin methylesterases (PMEs) (grouped within the pectin metabolism genes, PEC M, in Fig. 2d). These observations are consistent with previous reports on the regulation of cell wall genes by PIFs in darkness (Martin *et al*., 2016; Zhang *et al*., 2018; Zhang *et al*., 2021) and in shade conditions (Luo *et al*., 2022; Sénéchal *et al*., 2024). Moreover, some of these DEGs, such as XXT2 (Aryal *et al*., 2020), TSD2 (Krupková et al., 2007), PME2 (Hocq *et al*., 2024), or XTH19 (Miedes *et al*., 2013) have been previously implicated as key players for hook development and/or hypocotyl elongation in skotomorphogenesis. A similar, though less pronounced, downregulation of cell wall organization/biogenesis genes was also observed in *pifq* hypocotyls (Table S1, Fig. S6a), suggesting that PIFs regulate a specific subset of cell wall components in hook cells. Interestingly, recent evidence shows that alterations in cell wall mechanics caused by localized growth or external constraints might activate turgor-related signaling pathways that feed into a PIF4-depending network, modulating gene expression and cell elongation (Lorrai *et al*., 2024). Taken together, our results raise the possibility of a feedback loop between PIFs and the cell wall that contributes to balanced growth in apical hooks.

Our RNA-seq re-analysis also revealed a significant enrichment of microtubule motor activity (GO:0003777) among DEGs in *pifq* hooks (Table S1), with a striking proportion (32 of 58) of associated genes (mostly kinesins) upregulated and only one downregulated (Fig. 2e). Kinesins function contributes to the formation, maintenance, and function of cytoskeletal arrays and as drivers of long-distance transport of various cellular components (Nebenführ & Dixit, 2018). Several kinesins affect the spatial organization of cortical microtubules through a variety of mechanisms, such as cross- linking (Tian *et al*., 2015), polymerization and stabilization (Moschou *et al*., 2016), destabilization through cMT depolymerization (Oda and Fukuda, 2013), or reorientation by an unknown mechanism (Lan *et al*., 2023). It is likely that PIF-mediated repression of kinesins contributes to cMT reorganization and modulates differential growth during hook opening. The weaker misregulation of kinesin genes observed in *pifq* hypocotyls (Table S1, Fig. S6b) compared to *pifq* hooks (Fig. 2e) may indicate that PIFs play a more specific role in regulating cMTs homeostasis within the hook region.

In summary, this work demonstrates that PIFs are essential regulators of apical hook opening in dark-grown Arabidopsis. We propose that PIFs orchestrate cortical microtubule rearrangement (potentially through kinesin regulation) and cell expansion (likely via cell wall remodeling) to establish the differential growth patterns necessary for successful hook development. Further identification of the specific kinesins and cell wall-modifying activities involved will be key to fully elucidating the mechanistic basis of this developmental process

## MATERIAL AND METHODS

### Plant materials and growth conditions

*Arabidopsis thaliana* Col-0 was used as a wild-type. We introduced the cMTs reporter line p35S::GFP-MBD (Marc *et al*., 1998) into the *pifq* mutants (Leivar *et al*; 2008) by genetic crosses.

Seeds were surface sterilized for 2 min in 96% EtOH, transferred to 20% v/v NaOCl and 0.05% v/v Triton X-100 for 10 min, and rinsed five times with sterile distilled water. Sterilized seeds were sown on 0.5 MS (0.5 Murashige-Skoog basal salts, pH 5.8 and 0.8% w/v agar) in petri dishes and incubated at 4°C in darkness during 4-5 days for stratification. Chilled seeds were exposed to 3 h of white light in a growth chamber at 22°C to synchronize germination, and then kept in darkness in the same growth chamber for the indicated time points.

### Hypocotyl elongation and hook angle measurements

For hypocotyl elongation and hook angle measurements, seedlings were arranged horizontally on a plate, photographed using a digital camera and measured with ImageJ/Fiji software v1.53t. Hook angle was measured as the angle between the hypocotyl and an imaginary line between the cotyledons.

### Confocal microscopy and microtubule orientation measurements

Etiolated seedlings expressing p35S::GFP-MBD were imaged in a Carl Zeiss LSM 880 Indimo Axio Observer 7 inverted microscope using a dry objective C-Apochromat 40x/1.2 for cMTs orientation measurements; with the following settings to detect GFP signal: Excitation laser, 488 nm (2.4%); emission, 548 nm; gain, 650. The image size was 2140 × 2140 pixels and the pixel size: 0.1 μm x 0.1 μm x 1.0 μm. Z-stacks of the apical hook and single images of top half/upper-hypocotyl from the most external periclinal wall of epidermal cells were acquired. All the seedlings were mounted in water for image acquisition.

The analysis of cMTs distribution pattern was studied by fluorescence confocal microscopy in epidermal cells. The orientation and anisotropy of the cMTs was determined by the angle formed with the longitudinal axis of the cells using the FibrilTool plug-in designed for ImageJ/Fiji (Boudaoud *et al*., 2014), and categorized for each cell into transverse (0°-30°), oblique (30°-60°), and longitudinal (60°-90°) and random (Arico *et al*., 2024). For simplicity, only data from cMTs with transverse and longitudinal disposition are shown. Those data with anisotropy values below 0.05 were considered to be random (Arico *et al*., 2024). For each genotype, absolute frequency was calculated per category. At least three plants of each genotype were imaged per experiment and each experiment was repeated at least five times (total n° of cells=110- 204).

### Cell elongation measurements

Dark-grown seedlings were stained with Propidium iodide (PI) solution (1 mg/mL) mounted in 60 μl of water for image acquisition. Images to analyze cell length were taken with a Olympus IX83 inverted Disk Spinning Unit microscope with a Hamamatsu ORCA-Flash4.0 camera, using a dry objective UPLSAPO 20x/0.75. To detect RFP- DSU signals it used an emission laser, 585 nm with different exposure times according to the PI stain efficiency in each plant (in general 3-7s) and a lamp intensity of 6,00 V. The image size was adjusted in each image according to the plant size making up a mosaic, therefore the pixel size also varied. Z-stacks of the apical hook and single images of top half/upper-hypocotyl from the most external periclinal wall of epidermal cells were acquired. Cell wall contour determined by PI stain allows cell length measurements using the segmented line tool of ImageJ/Fiji on mosaic images of each plant. Cells were classified in concave and convex sides of the apical hook, and the following 4 up to 6 cells on the hypocotyl were considered as apical hypocotyl zones. At least three plants of each genotype were imaged per experiment and each experiment was repeated at least five times (total n° of cells=60-270).

### Scanning electron microscopy

Two- and three-day dark-grown p35S::GFP-MBD in wild-type and *pifq* background samples were fixed in FAA solution (3.7 % formaldehyde, 5 % glacial acetic acid, 50 % ethanol), and washed with PBS 48 h later. After dehydration with a series of ethanol solutions (50%, 60%, 70%, 80%, 90%, 96% for 40 min and 100% for 12 hours), they were transferred to pure acetone for desiccation using CO2 in an automated critical point dryer (EM CPD300, Leica). Finally, samples were observed and photographed using a FEI QUANTA 200 field emission gun scanning electron microscope (SEM) located at the Centro Científico Tecnológico CONICET Rosario (Argentina). The observations were conducted in low vacuum mode (0.8 mBar) at 5 Kv and 10 mm working distance.

### Statistical analyses

Data were analyzed using Student’s t-test when two sample datasets were compared or one-way ANOVA for more than two datasets, and the differences between means were evaluated using Tukey’s post-test (Infostat Software v.2020I, Facultad de Ciencias Agropecuarias, Universidad Nacional de Córdoba, Cordoba, Argentina). Statistically significant differences were defined as those with a P-value <0.05 (*), <0.01 (**), <0.001 (***), or with different letters (for one-way ANOVA, Tukey post-test) as indicated.

## Acknowledgements

We thank Diego Aguirre for plant care in growth chambers; Pablo Pomata, Sebastian Petracca and Rodrigo Vena for technical assistance with confocal microscopy. This research was funded by CONICET (PIP1985 to MAM and N G-S).

## Author contributions

NB, PV, FS, MAM, and NG-S performed the experiments; NB, PV, MAM, and NG-S analyzed the data; MAM, and NG-S conceived the project and wrote the manuscript; MAM, and NG-S provided the funding and revised the manuscript. All authors read and approved the final manuscript.

## Supplemental material

Figure S1. Two-step dynamics of apical hook opening.

Figure S2. Hypocotyl and cell growth dynamics of p35S:GFP:MBD lines. Figure S3. Cell growth dynamics of p35S:GFP:MBD *pifq* hooks.

Figure S4. Hypocotyl and cell growth dynamics of p35S:GFP:MBD *pifq* lines.

Figure S5. Scanning electron microscopy (SEM) images within the apical hooks of WT and *pifq* mutants.

Figure S6. Cell wall genes are misregulated in *pifq* hypocotyls. Table S1. RNA-seq re-analysis.

