## Supplemental Material for "PHYTOCHROME INTERACTING FACTORS are required to coordinate microtubule dynamics and differential cell growth during Arabidopsis apical hook opening"

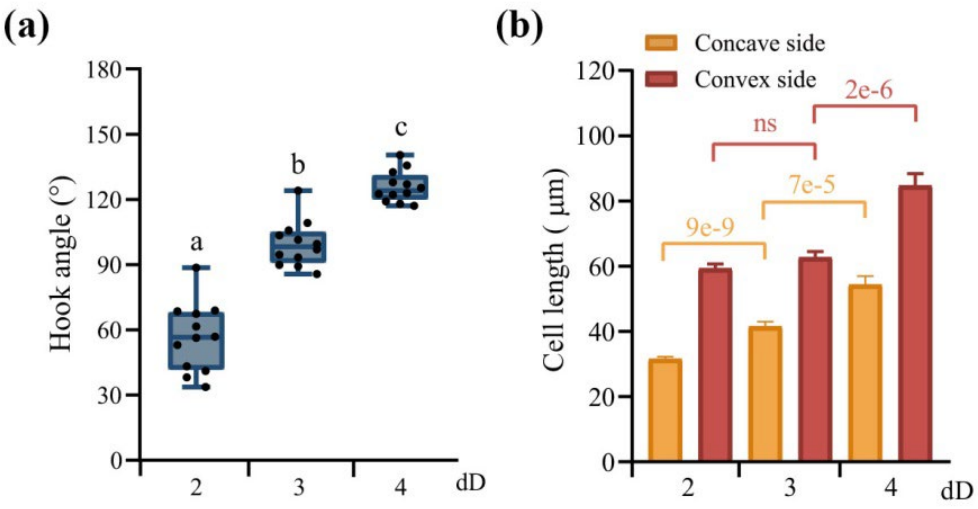


**Figure S1. Two-step dynamics of apical hook opening.** (a) Hook angle measurements of dark-grown p35S:GFP:MBD at indicated time points. dD indicates days in the dark. Data are reported as a box plot with median (horizontal line inside the box), and points represent biological replicates of 30-52 seedlings each. Statistically significant differences between groups are indicated by different letters (one-way ANOVA, $P < 0.05$). Associated data with hook opening rate between time points is presented in Fig. 1c. (b) Cell length measurements on the concave and convex sides of dark-grown p35S:GFP:MBD apical hooks at indicated time points. Data are reported as means ± SEM of at least five biological replicates (total n° of cells=106-189). Statistical significance between samples is indicated by p-values (t-test; ns, not significant). Associated data with cell growth rates between time points are presented in Fig. 1d.

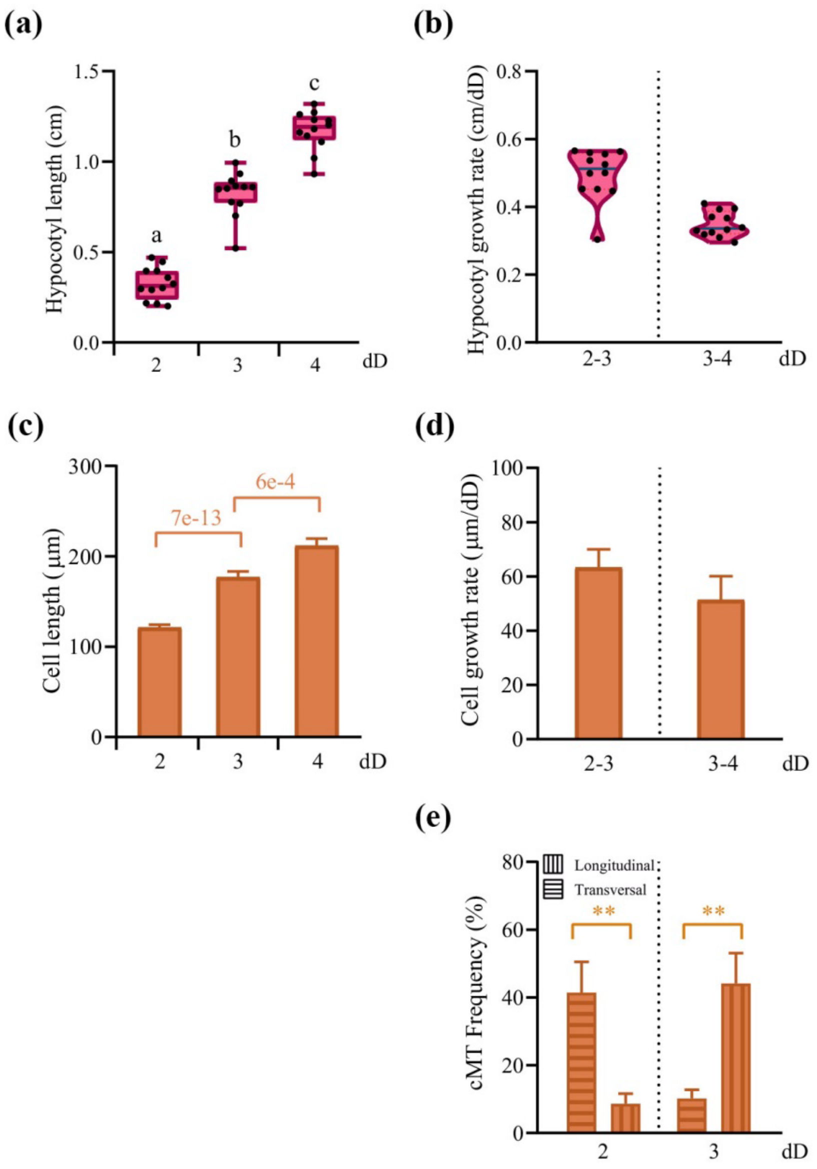


**Figure S2. Hypocotyl and cell growth dynamics of p35S:GFP:MBD lines.** (a) Hypocotyl length measurements of dark-grown p35S:GFP:MBD at indicated time points. dD indicates days in the dark. Data are reported as a box plot with median (horizontal line inside the box), and points represent biological replicates of 30-52 seedlings each. Statistically significant differences between groups are indicated by different letters (one-way ANOVA, $P < 0.05$). (b) Hypocotyl growth rate associated with data in (a). Data are reported as violin plots with median in blue. (c) Cell length measurements on hypocotyls of dark-grown p35S:GFP:MBD at indicated time points. Data are reported as means ± SEM of six biological replicates (total n° of cells=183-254). Statistical significance between samples is indicated by p-values (t-test; ns, not significant). (d) Cell growth rate associated with data in (c). Data are reported as means ± SEM. (e) Frequency of longitudinal and transversal cMTs in p35S::GFP-MBD hypocotyls at indicated time points. Data are reported as means ± SEM of at least nine biological replicates (total n° of cells=148-204). Asterisks indicate statistically different mean values (t-test, **, $P < 0.01$).

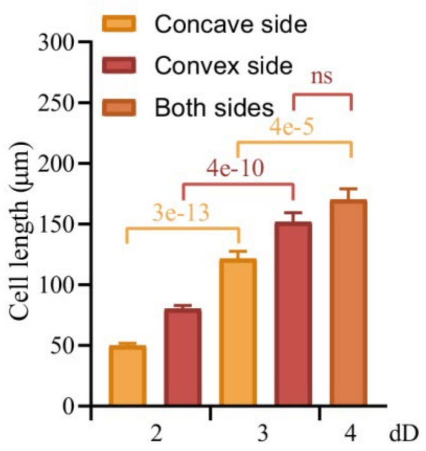


**Figure S3. Cell growth dynamics of p35S:GFP:MBD *pifq* hooks.** Cell length measurements on the concave and convex sides of dark-grown p35S:GFP:MBD *pifq* apical hooks at indicated time points. dD indicates days in the dark. Data are reported as means ± SEM of at least five biological replicates (total n° of cells=60-86). Statistical significance between samples is indicated by p-values (t-test; ns, not significant). Associated data with cell growth rates between time points are presented in Fig. 2b.

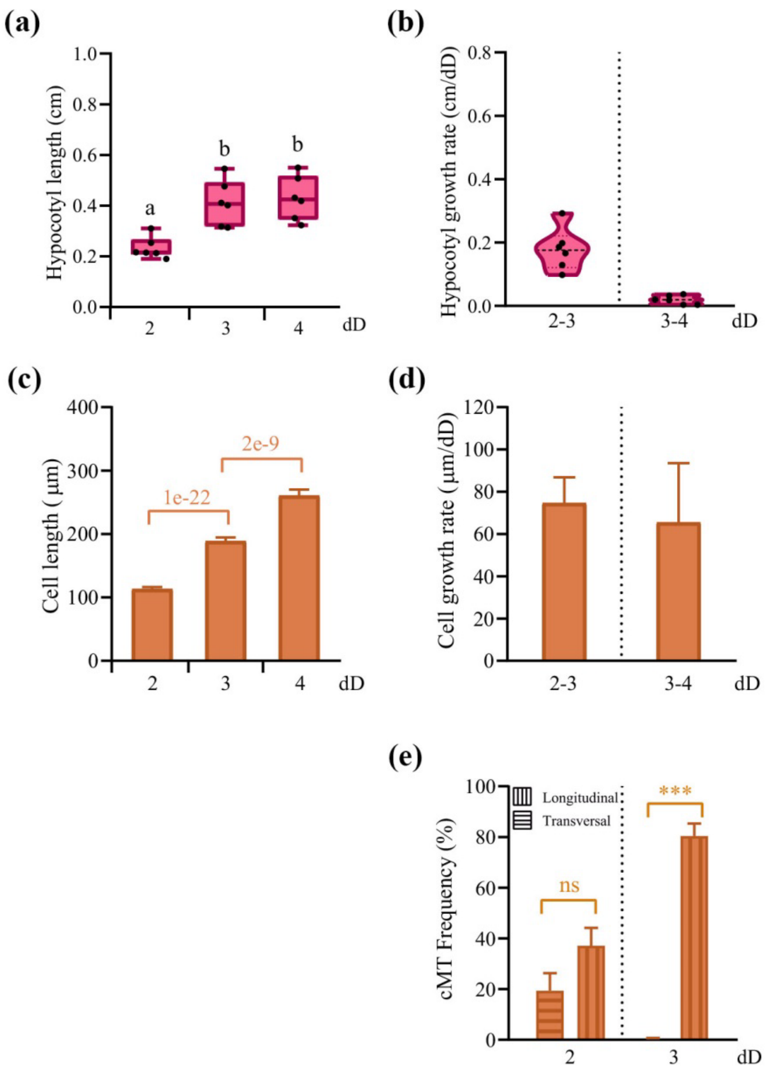


**Figure S4. Hypocotyl and cell growth dynamics of p35S:GFP:MBD *pifq* lines.** (a) Hypocotyl length measurements of dark-grown p35S:GFP:MBD *pifq* at indicated time points. dD indicates days in the dark. Data are reported as a box plot with median (horizontal line inside the box), and points represent biological replicates of 30-53 seedlings each. Statistically significant differences between groups are indicated by different letters (one-way ANOVA, $P < 0.05$). (b) Hypocotyl growth rate associated with data in (a). Data are reported as violin plots with median in black (dashed line). (c) Cell length measurements on hypocotyls of dark-grown p35S:GFP:MBD *pifq* at indicated time points. Data are reported as means ± SEM of six biological replicates (total n° of cells=195-270). Statistical significance between samples is indicated by p-values (t-test; ns, not significant). (d) Cell growth rate associated with data in (c). Data are reported as means ± SEM. (e) Frequency of longitudinal and transversal cMTs in p35S::GFP-MBD *pifq* hypocotyls at indicated time points. Data are reported as means ± SEM of at least seven biological replicates (total n° of cells=110-147). Asterisks indicate statistically different mean values (t-test, ***, $P < 0.001$).

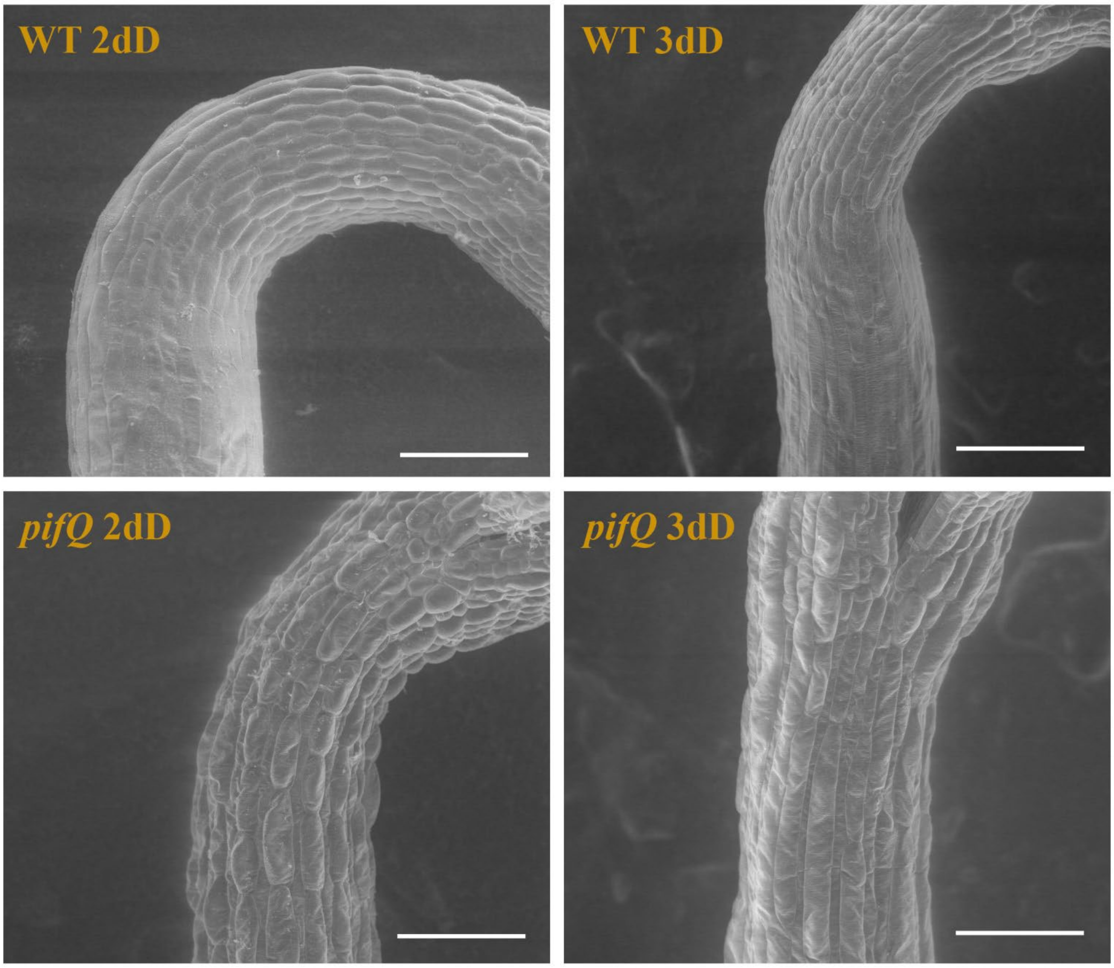


**Figure S5. Scanning electron microscopy (SEM) images within the apical hooks of WT and *pifq* mutants.** Seedlings were grown in darkness for two or three days (bar=100 μm).

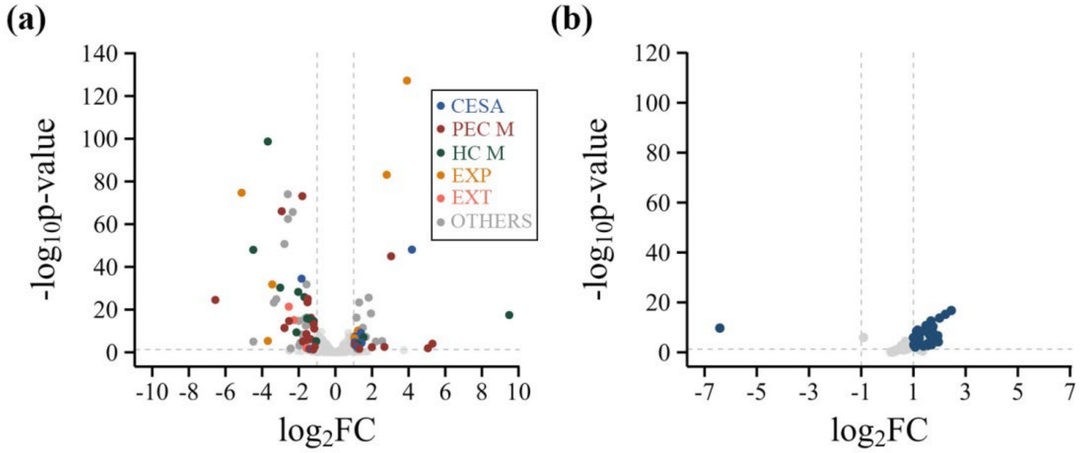


**Figure S6. Cell wall genes are misregulated in *pifq* hypocotyls.** (a,b) Volcano plots showing log2FC (*pifq*/WT) vs –log10p-value for genes belonging to cell wall organization or biogenesis (GO:0071554) (a), or microtubule motor activity (GO:0003777) (b) from four-day hypocotyls dark-grown *pifq* and WT RNA-seq re-analysis (Zhang *et al*., 2021). Associated values are presented in Table S1. Horizontal dashed lines represent the p-value threshold, while vertical dashed lines represent cut-off expression values. CESA: cellulose synthase genes; PEC M: pectin metabolism associated genes; HC M: Hemicellulose metabolism associated genes; EXP: expansins; EXT: extensins.
